# *Cocculus hirsutus*-derived phytopharmaceutical drug has potent anti-dengue activity

**DOI:** 10.1101/2020.09.18.303149

**Authors:** Ankur Poddar, Rahul Shukla, Hemalatha Beesetti, Upasana Arora, Ravi Kant Rajpoot, Rajgokul K Shanmugam, Srinivas Palla, Kaushal Nayyar, Deepika Singh, Venugopal Singamaneni, Prasoon Gupta, Ajai Prakash Gupta, Sumeet Gairola, Y. S. Bedi, Tapesh Jain, Bhupendra Vashishta, Ravindra Patil, Harish Madan, Sumit Madan, Rinku Kalra, Ruchi Sood, Ram Vishwakarma, Altaf A Lal, Navin Khanna

## Abstract

**Background:** Dengue is a serious public health concern worldwide, with ~3 billion people at risk of contracting dengue virus (DENV) infections. Currently, no effective vaccine or drug is available for the prevention or treatment of dengue, which leaves only anti-mosquito strategies to combat this disease. The present study was initiated to determine the *in-vitro* and *in vivo* protective effects of a plant-derived phytopharmaceutical drug for the treatment of dengue.

**Methodology/Principal Findings:** In our previous report, we had identified methanolic extract of the aerial parts of *Cissampelos pareira* to exhibit *in vitro* and *in vivo* anti-dengue activity against all the four DENV serotypes. In the current study, we have identified another Indian medicinal plant, *Cocculus hirsutus*, which has a more potent anti-dengue activity than *C. pareira.* The activity has been evaluated through flow-cytometry-based virus inhibition assay. Interestingly, the stem of *C. hirsutus* was found to be more potent than the aerial part irrespective of the extraction solvent used viz., denatured spirit, hydro-alcohol (50:50) and water. Hence, the aqueous extract of stem of *C. hirsutus* (AQCH) was further advanced for investigations because of greater regulatory acceptance. The AQCH exhibited dose-dependent inhibition of release of DENV and its secretory antigen, NS1. Five chemical markers viz. Sinococuline, 20-Hydroxyecdysone, Makisterone-A, Magnoflorine and Coniferyl alcohol were identified as the major chemical ingredients of the AQCH extract. These chemicals were subsequently used for extract standardisation. Importantly, AQCH completely protected AG129 mice at 25 mg/kg/dose body weight when fed 4 times a day post-infection with a lethal dose of DENV-2 S221 strain. Because of its potential as an effective phytopharmaceutical drug against dengue, AQCH, has been formulated into tablets for further pre-clinical and clinical developments.

**Conclusions/Significance:** We provide evidence of the pan anti-dengue potential of *C. hirsutus-*based phytopharmaceutical drug as determined through *in vitro* and *in vivo* experiments. We have also characterized five chemical entities in the drug substance, which provides means for standardization of drug substance and drug product. Based on these findings, a program to develop a safe and effective *C. hirsutus-*derived phytopharmaceutical drug for the treatment of dengue has been initiated.

**Author summary:** There is an urgent need to develop a safe and effective drug against dengue, which is a rapidly expanding mosquito-borne viral disease. Half of the world’s population has been estimated to be at risk of contracting this disease and the situation remains grim due to lack of an approved drug. We aimed to develop an ethnopharmacological drug against dengue by exploring traditional Indian medicinal science, Ayurveda. This led us to identify a creeper, *Cocculus hirsutus*, as a more potent anti-dengue plant than *Cissampelos pareira,* reported in our earlier published study. The stem part of *C. hirsutus* was found to be more efficacious in inhibiting the propagation of dengue viruses (DENVs) in cell culture than its aerial part. Hence, we chose to advance aqueous extract of stem of *C. hirsutus* (AQCH) for further studies. Importantly, AQCH also protected immune-compromised mice from lethal DENV infection, which is suggestive of its potential clinical relevance. We have identified five chemical marker compounds in AQCH to gauge the quality and consistency of extract preparation and its formulation into stable tablets. Based on the findings of this study, we have undertaken the development of a safe and effective *C. hirsutus-*derived phytopharmaceutical drug for the treatment of dengue.

## Introduction

Dengue is a mosquito-borne disease caused by infection of any of the four antigenically distinct dengue virus (DENV) serotypes, which belong to genus *Flavivirus* of *Flaviviridae* family of positive single stranded RNA viruses. Dengue infection has the potential of causing a pandemic with increased outbreaks in many parts of the world [1]. Climate change, population growth, increased international travel, rapid urbanization and ineffective vector control strategies have led to the expansion of dengue footprint worldwide. Dengue is endemic in more than 100 countries of South-East Asia, Eastern Mediterranean, Western Pacific, Americas and Africa [2–4]. The co-circulation of multiple DENV serotypes leading to concurrent infections have also been reported in hyper-endemic nations [5,6].

A recent study estimated that every year around 390 million dengue infection cases take place and of these 96 million result in clinical manifestations [7]. Another study revealed that around 3.9 billion people are at risk of contracting dengue disease, making it a serious global health concern [8]. Asia alone is saddled with 70% of the global dengue burden and India is hyperendemic for dengue with a whopping 34% contribution to global burden [7]. It has been estimated that 49% of India’s population has already been infected with the DENV, however, the prevalence varies among different regions and age groups [9]. According to a Global Burden of Disease study, dengue alone inflicts a global burden of USD 8.9 billion as per 2013 prices [10].

Dengue is transmitted among humans through the bite of infected female mosquitoes, primarily *Aedes aegypti*. This results in asymptomatic dengue infection in majority of the infected population, however, 20-25% of infected individuals develop symptomatic disease that persists for 2-7 days post 4-7 days of incubation after the mosquito bite [4,11].

Symptomatic dengue infection can range from uncomplicated dengue without warning signs like fever, rash, retro-orbital pain, etc., to dengue with warning signs of fluid accumulation, mucosal bleeding, etc., and severe dengue characterized by severe plasma leakage, severe bleeding, respiratory distress and organ deterioration [4,11,12].

As of today, no specific treatment is available for dengue and patients are provided only supportive medical care, especially fluid management [11]. The recent launch of a dengue vaccine, Dengvaxia, by Sanofi Pasteur is of limited use due to its disease enhancement concerns in seronegative vaccinees [13–15].

Though several attempts have been made for development of drugs against dengue, the efforts have not yielded a safe and effective drug so far. [16–18]. Several drugs such as chloroquine [19], celgosivir [20], lovastatin [21], balapiravir [22], prednisolone [23] have been evaluated for their ability to treat dengue infection and disease. All these trials failed to meet the efficacy endpoints.

A parallel effort of investigating the plant extracts widely known for their traditional use by tribals, traditional healers and local people against dengue-like febrile illnesses has been undertaken [24]. Some of these plants are *Azadirachta indica* [25], *Hippophae rhamnoide* [26], *Carica papaya* [27], *Cissampelos pareira* [28], etc. These herbal formulations, if validated and proved for their clinical efficacy, can form basis for the development of safe and effective drugs against dengue.

In our previous study, we identified *C. pareira* plant for its anti-dengue activity against all four DENV serotypes [28]. In the current study, we have expanded the investigation by further exploring herbal repertoire to identify a plant that may have more potent anti-dengue activity and could become promising candidate of dengue drug development program.

We have identified *Cocculus hirsutus,* which belongs to the class Magnoliopsida and family Menispermaceae, to have more potent anti-dengue activity than *C. pareira*. This plant is traditionally known to possess many medical properties as it is used as a detoxifier, aphrodisiac, antipyretic, diuretic, laxative, cardiotonic, chronic rheumatism, syphilitic cachexia, skin diseases, constipation, kidney problems, etc [29]. The acute toxicity of aqueous extract of aerial parts of *C. hirsutus* has been established to be >3000 mg/kg body weight in Swiss mice [30].

Our study reports for the first time the anti-dengue potential of *C. hirsutus* based on exhaustive scientific validations. We have examined *C. hirsutus* by preparing extracts of its aerial and stem parts using different solvents and varied drying conditions. For all the extracts prepared, anti-dengue activity was established through a flow-cytometry-based virus inhibition assay. Purified aqueous extract of stem of *C. hirsutus* (AQCH) was found to possess the highest pan-DENV inhibitory activity. AQCH demonstrated its ability to reduce NS1 and virus secretion in the supernatant in an *in vitro* analysis in a dose-dependent manner. Through chemical fingerprinting analyses, five marker compounds viz.

Sinococuline, Magnoflorine, 20-Hydroxyecdysone, Makisterone-A and Coniferyl alcohol, were identified to be present consistently in multiple AQCH batches. The AQCH also protected the AG129 mice when challenged with lethal dose of DENV-2 S221 strain, which further augments its potential for development as a phytopharmaceutical drug against dengue.

## Methods

### Animal ethics statement

This study involved experiments on AG129 mice, which were performed at the International Centre for Genetic Engineering and Biotechnology, New Delhi (ICGEB/IAEC/08/2016/RGP-15) in compliance with the ‘Committee for the Purpose of Control and Supervision of Experiments on Animals’ guidelines issued by the Government of India.

### Cells and viruses for *in vitro* and *in vivo* DENV inhibition assays

Vero cell line was purchased from the American Type Cell Culture (ATCC), Virginia, USA. This monkey kidney cell line was maintained using Dulbecco’s Modified Eagle medium (DMEM) supplemented with 10% ΔFBS, in a 10% CO_2_ humidified incubator at 37°C. WHO reference DENV strains DENV-1 (WP 74), DENV-2 (S16803), DENV-3 (CH53489), and DENV-4 (TVP-360) were received from Dr. Aravinda de Silva’s lab, University of North Carolina (UNC), USA. These viruses were cultured in C6/36 cells procured from ATCC, Virginia, USA. Mouse adapted DENV-2 S221, used in animal experiments, was procured from Global Vaccines Inc., North Carolina, USA and cultured in DMEM adapted C6/36 cells. Dengue cross-reactive monoclonal antibodies (mAbs), 4G2 and 2H2, which recognize epitopes in fusion loop and prM protein of DENVs respectively, were produced in-house from their respective hybridomas procured from ATCC, Virginia, USA. 2H2 mAb was labelled in-house with Alexa fluor-488 through commercial labelling kit (Thermo Fischer Scientific, Eugene, USA).

### Chemicals and reference compounds for HPLC chromatography

Analytical or HPLC grade organic solvents used in the plant extraction and HPLC analysis, were procured from E. Merck Ltd., Mumbai, India. Prior to use, solvents were filtered through a 0.45 μm membrane filter (Millipore, Billerica, MA, USA). The HPLC column RP18e Purospher-STAR (Hibar) (250 × 4.6 mm; 5 μm) was used for chemical fingerprinting (E. Merck) and Eclipse 5 μm column (9.4 × 250 mm) was used for the purification of marker compounds (Agilent). The chemicals and reagents used for standardisation and quality control were procured from Sigma-Aldrich, USA. Water for extraction and HPLC analysis was obtained from high-purity Milli-Q Advantage A10 water purification system (Millipore, Molsheim, France).

### Plant procurement, validation, and extract preparation

The botanical raw materials (BRMs), i.e., aerial or stem parts of *C. pareira* and *C. hirsutus,* were collected by the botanist. Identification of the collected BRMs was performed at the Plant Science Division of CSIR-Indian Institute of Integrative Medicine (CSIR-IIIM), Jammu, India. Duly identified herbarium specimens of *C. pariera* (Accession No. RRLH-23148) and *C. hirsutus* (Accession No. RRLH-23152) were submitted to the internationally recognized Janaki Ammal Herbarium (RRLH) at CSIR-IIIM, Jammu. Further, for some studies, BRM of *C. hirsutus* collected from the same area was procured from the local vendor. After critical macroscopic and microscopic examinations, the botanical identity of the procured BRM samples of *C. hirsutus* were confirmed at the Plant Science Division of CSIR-IIIM. The duly identified samples of the procured BRMs, i.e., aerial (Accession Nos. CDR-4037, and CDR-4038) and stem (Accession Nos. CDR-4061, CDR-4064, CDR-4065, and CDR-4078) parts of *C. hirsutus* have been submitted to the Crude Drug Repository (CDR) at CSIR-IIIM, Jammu.

Post-confirmation of botanical identity of BRM, extracts were prepared in the extraction solvent (denatured spirit, hydro-alcohol 50:50 and water).

### Flow-cytometry-based virus inhibition assay

Vero cells were seeded in a 96-well plate (20,000-25,000 cells/well) in 200 μl DMEM + 10% ΔFBS and incubated for 24 hr in an incubator adjusted at 37°C and 10% CO_2_. Next day, cells were infected with 100 μl of DENV-1, −2, −3, and −4 dilutions, prepared in DMEM + 0.5% ΔFBS (dilution media), to yield ~10% infection. After a 2 hr incubation of Vero cells with the virus at 37°C, 10% CO_2_, virus was aspirated and 200 μl of a suitable range of AQCH prepared in dilution media was added to the wells in duplicates. Cells were incubated further for another 46 hr in an incubator at 37°C, 10% CO_2_. Wells infected with the virus but without any subsequent extract treatment served as virus controls whereas wells with no infection and no treatment served as cell controls. These experimental controls were utilized for relative % virus infection calculations and antibody background signal adjustments, respectively. After completion of the incubation period, cells were stained for the presence of cytosolic DENVs with Alexa-488 labelled 2H2 mAb. For staining, media was aspirated from the top of the cells and washed with 150 μl PBS. Cells were trypsinised and transferred to a 96 well U bottom plate. After transfer, cells were centrifuged at 1500 rpm for 5 mins and supernatant was aspirated. Cells were washed with PBS again and then fixed with 50 μl 4% para-formaldehyde for 20 mins. Cells were centrifuged at 2500 rpm for 5 mins and supernatant was aspirated. Cells were washed twice with 150 μl permeabilization or perm buffer and blocked with 40 μl 1% normal mouse sera (prepared in perm buffer) for 30 mins. Without removing the blocking solution, 20 μl Alexa-488 labelled 2H2 mAb (prepared in blocking solution) was added to stain the cells for DENVs and incubated for 1 hr at 37°C with gentle shaking. Post-incubation, cells were centrifuged at 2500 rpm for 5 mins and supernatant aspirated. Cells were washed twice with perm buffer and re-suspended in 100 μl of PBS. The above processed cells were analysed through a BD FACS Verse flow cytometer and 5000 cells were counted per well. Data was analyzed through FlowJo software to determine the relative percentage of infected cells for each test substance concentration with respect to virus only control group. The 50% inhibitory concentration (IC_50_) of the test substance was determined as the concentration that inhibited 50% of dengue virus infection with respect to virus control, calculated using non-linear regression analysis of GraphPad Prism software.

### NS1 ELISA assay

Vero cells were seeded in a 48 well plate (40,000 cells/well) in 500 μl DMEM + 10% ΔFBS and incubated at 37°C, 10% CO_2_. Next day, cells were infected with DENV-1, −2, −3, and −4 at 0.1 MOI prepared in dilution media. After 2 hr of infection, media was aspirated and the cell monolayer was overlaid with 200 μl of different concentrations of extract (100, 50, 25, and 12.5 μg/ml) prepared in dilution media. Wells that did not receive any infection but only dilution media served as negative control. Every day 10 μl of overlaying culture supernatant was withdrawn from each well for 6 days post-infection for the detection of NS1 antigen using commercial Dengue NS1 Ag Microlisa kits (J. Mitra & Co. Pvt. Ltd.).

### Extracellular viral estimation

In this assay, culture supernatant of DENV-infected cells were collected and the virus was titrated in both the AQCH treated and untreated wells. Briefly, Vero cells were seeded and infected with DENVs in six 96 well plates as detailed in flow-cytometry-based virus inhibition assay. After 2 hr of infection, the virus infection media was aspirated and the monolayer was overlaid with 200 μl of different concentrations of AQCH (100, 50, 25, and 12.5 μg/ml) prepared in dilution media. Post 24 hr of infection, one plate was harvested each day for the next 6 days and the supernatant was transferred to a 96-well U-bottom plate. Collected samples were stored at 4°C till Day 6. The titre of DENVs in these collected supernatants was evaluated on Vero cells in flow-cytometry-based assay [31] to yield FACS Infectious Units per ml (FIUs/ml).

### MTT assay

The *in vitro* cell cytotoxic index (CC_50_) of AQCH was evaluated through MTT (3-(4, 5-dimethylthiazolyl-2)-2, 5- diphenyltetrazolium bromide) assay. Vero cells were seeded as described in flow-cytometry based virus inhibition assay. Post 24 hr incubation at 37°C and 10% CO_2_, overlay media was removed and 200 μl of a suitable concentration range of AQCH prepared in dilution media was added to the wells in duplicates; cells incubated with dilution media alone were kept as cell control and processed in parallel. Cells were incubated further for another 46 hr in an incubator at 37°C, 10% CO_2_. Post incubation, 10 μl of 5 mg/ml MTT reagent (procured from Sigma Aldrich, USA) prepared in PBS was added and further incubated for 2 hr at 37°C, 10% CO_2_. Upon formation of formazan crystals, the overlay was removed and 100 μl of DMSO was added. After the dissolution of crystals in DMSO, absorbance was taken at 570 nm. The % cell cytotoxicity was calculated for each AQCH concentration with respect to cell control. The concentration of AQCH at which 50% cell cytotoxicity was observed is reported as CC_50_.

### AQCH chemical fingerprinting and characterization

AQCH was characterized by chemical fingerprinting using RP18e Purospher-STAR (Hibar) (250 × 4.6 mm; 5 μm) column. The mobile phase containing a buffer (0.1% formic acid in water) and acetonitrile was used at a flow rate of 0.65 ml/min at a column temperature of 30°C and monitored at 254 nm. The compounds were isolated by column chromatography using silica gel (60-120 and 230-400 mesh); fractions were monitored by TLC using pre-coated silica gel plates 60 F254 (Merck) and spots were visualized by UV light or by spraying with H_2_SO_4_-MeOH, anisaldehyde-H_2_SO_4_ reagents. The isolated compounds were characterized by NMR and mass spectrometry using Bruker 400 MHz spectrometer and Agilent 1100 LC-Q-TOF, respectively.

### *In vivo* evaluation of the efficacy of AQCH in AG129 primary dengue lethal mouse model

AG129 mice deficient in IFN-α/β and IFN-γ receptors were purchased from the B&K Universal, United Kingdom, and housed and bred at the International Centre for Genetic Engineering and Biotechnology (ICGEB), New Delhi. Experimental mice (six per group), 6 to 8 weeks old, were housed in BSL-2 containment facility. They were intravenously injected with a lethal dose (1.0 × 10^5^ FIU) of mouse-adapted DENV-2 strain S221. Half an hour post DENV-2 S221 infection, mice were fed orally four times a day (QID) with either 8.25 or 25 mg/kg/dose of AQCH for a period of five days. Three groups of mice served as experimental controls. First, ‘Uninfected’, that was neither infected with DENV-2 S221 nor was treated with AQCH. Second, ‘Only AQCH’, which was not infected with DENV-2 S221 but treated with AQCH. Third, ‘V’ that was infected with DENV-2 S221 but was not treated with AQCH. All mice groups were monitored for their survival, body weight change and morbidity score for a period of 15 days post-infection. Statistical evaluation of survival score was performed through Log Rank (Mantel Cox) test and p value <0.05 was considered statistically significant. Morbidity score was based on 5 point system: 0.5, mild ruffled fur; 1.0, ruffled fur; 1.5, compromised eyes; 2, compromised eyes with hunched back; 2.5, loose stools; 3.0, limited movement; 3.5, no movement/hind leg paralysis; 4.0, euthanized if cumulative score was 5. All AQCH doses used for feeding were prepared at once in water with 0.1% methylcellulose (v/v) and stored at 4°C. An appropriate volume of doses was pre-incubated at room temperature before feeding to animals. For a QID dosing, mice were fed with AQCH at 4/4/4/12 hr cycle (7 AM, 11 AM, 3 PM and 7 PM on each day for 5 days).

### *In vitro* evaluation of the interaction of paracetamol with AQCH

Interaction between paracetamol and AQCH was determined *in vitro* through flow-cytometry-based virus inhibition assay. The 24 hr seed vero cells were infected for 2 hr with DENV-1 (as described in the flow-cytometry-based virus inhibition assay). Post incubation, virus infection media was aspirated and cells were treated for 46 hr with AQCH concentrations ranging from 0 to 25 μg/ml along with parallel treatment of 1, 10 and 100 μg/ml paracetamol in different lanes of a 96-well plate. Each treatment was evaluated in duplicates and IC_50_ was calculated through non-linear regression analysis using GraphPad Prism.

### Stability analyses of AQCH and its tablets

AQCH was formulated into tablets of different strengths (100 mg, 300 mg, and 500 mg) using the approved excipients. The accelerated and long-term stability of AQCH and AQCH tablets was assessed by exposing them to different conditions (30 ± 2°C/ 65 ± 5% RH, and 40 ± 2°C/ 75 ± 5% RH). The *in vitro* anti-dengue activity was evaluated for the stored samples at different time points (1, 2, 3, and 6 months) through flow-cytometry-based virus inhibition assay.

## Results

### Selection of *Cocculus hirsutus* for the evaluation of anti-dengue activity

Guided by the Indian Ayurveda literature and our *in vitro* and *in vivo* bioassays, we had earlier identified and established that methanolic extract of aerial parts of *C. pareira* possesses pan anti-dengue activity [28]. *C. pareira* belongs to family Menispermaceae, which is historically known to be rich in a variety of alkaloids [32]. Menispernaceae family is divided into eight tribes and three sub-tribes and consists of ~72 genera [32]. In our previous study [28], a total of 19 plants were evaluated for their anti-dengue activity, two of which, *C. pareira* and *Tinospora cordifolia*, belonged to the family Menispermaceae; both of them were found to possess anti-dengue activity. However, *C. pareira* which belongs to Cocculeae tribe of Menispermaceae exhibited significantly stronger anti-dengue activity than *T. cordifolia* which belongs to Tinosporeae tribe.

Thus, in our quest to find a more potent anti-dengue plant, we focussed our search on plants belonging to Cocculeae tribe of Menispermaceae. An indole alkaloid, hirsutine, derived from *Uncaria rhynchophylla*, was recently reported to inhibit later stages of DENV life cycle [33].

Amongst Menispermaceae, hirsutine alkaloids have largely been reported to be present in *C. hirsutus* which like *C. pareira*, belongs to Cocculeae tribe [32, 34]. Thus, it was decided to explore *C. hirsutus* for anti-dengue potential.

### *C. hirsutus* possesses pan anti-dengue activity and is more potent than *C. pareira*

With methanolic extract of aerial parts of *C. pareira* as our benchmark [28], we prepared three batches each of methanolic extract of aerial parts of *C. pareira* and *C. hirsutus*. In this study, we evaluated all these six methanolic extracts for anti-dengue activity in an *in vitro* flow-cytometry-based virus inhibition assay instead of conventional plaque based bioassay used previously [28]. The flow-cytometry-based virus inhibition and plaque based bioassays are principally similar. However, evaluations made through flow-cytometry-based virus inhibition assay are advantageous because it is high-throughput and is more stringent as it uses a higher dose of DENV for the evaluation of anti-dengue activity. In the flow-cytometry-based virus inhibition assay used in the current study, the Vero cells were infected with DENVs, and post-infection cells were incubated in media containing extract at various concentrations for 46 hr.

Post-incubation cells were fixed, permeabilized and stained with Alexa fluor labelled anti-dengue mAb, 2H2, reactive to all the four DENV serotypes, which were read in a flow cytometer to determine the percentage of DENV infected cells. This was used to calculate the extract concentration at which 50% of the DENV infection was inhibited (IC_50_). Upon parallel evaluation of all the six extracts in flow-cytometry-based virus inhibition assay, it was observed that all the three batches of methanolic aerial *C. hirsutus* extracts possessed significantly stronger anti-dengue activity against all the four DENV serotypes as compared to methanolic aerial *C. pareira* extracts (Fig 1).

**Fig 1:**
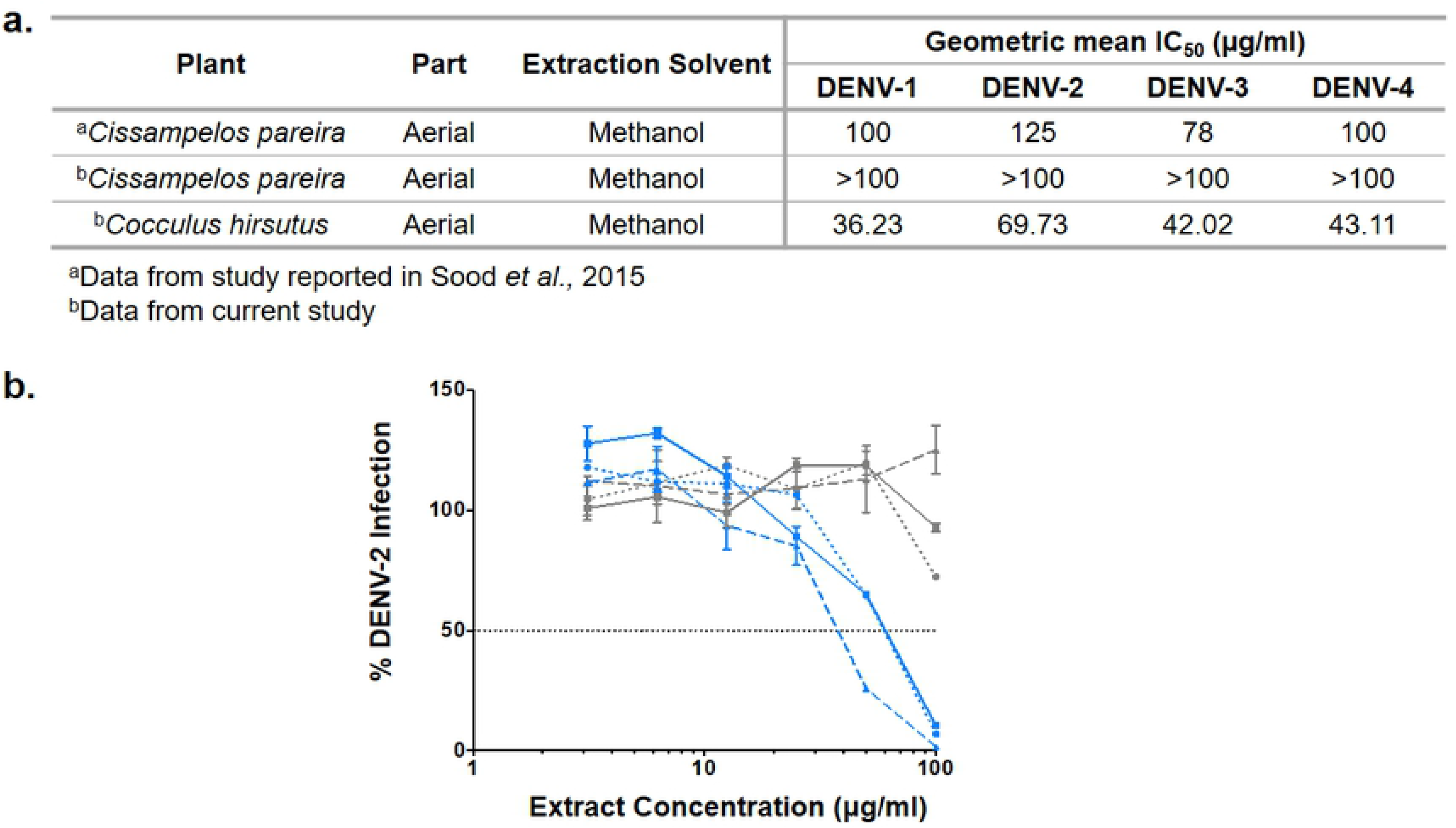
*C. hirsutus* possesses a more potent anti-dengue activity than *C. pareira*: Three batches each of the methanolic extracts of aerial parts of *C. pareira* and *C. hirsutus* were evaluated for their anti-dengue activity at 3.12, 6.25, 12.5, 25, 50 and 100 μg/ml extract concentrations, and the % DENV infection was recorded in a flow-cytometry-based virus inhibition assay against all the four DENV serotypes. The concentration of extract (μg/ml) that resulted in 50% inhibition of viral infection as compared to virus control was calculated as IC_50_ using Graphpad Prism. (a) IC_50_ values were calculated separately for each of the three extracts prepared from both the plants and their geometric mean IC_50_ values against each of the four DENV serotypes were calculated as reported in the table. IC_50_ values of aerial methanolic *C. pareira* extract from plaque reduction neutralisation assay reported in Sood *et al.*, 2015 and taken as reference for the current study are also shown in the table, (b) Graph of % DENV-2 infection observed with each of the six extracts with grey and blue curves representing the three *C. pareira* and *C. hirsutus* aerial methanolic extracts, respectively. Dashed horizontal line represents 50% DENV-2 infection value.

### Selection of aqueous extract of stem of *C. hirsutus* for further evaluation

Upon selection of *C. hirsutus* over *C. pareira* because of its more potent anti-dengue activity, we explored individual preparation of extracts of both the aerial (Fig 2a; dashed curves) and stem parts (Fig 2a; solid curves) of *C. hirsutus* in various solvents viz. denatured spirit, hydro-alcohol (50:50) and water. These extracts were evaluated against all the four DENV serotypes by flow-cytometry-based virus inhibition assay and their IC_50_ values were compared (Fig 2b). It was observed that the stem part of *C. hirsutus* was significantly more potent than aerial part irrespective of the extraction solvent used. Thus, aqueous extract of stem of *C. hirsutus*, hereafter referred to as AQCH, was advanced further due to simpler regulatory compliance associated with aqueous extracts.

**Fig 2:**
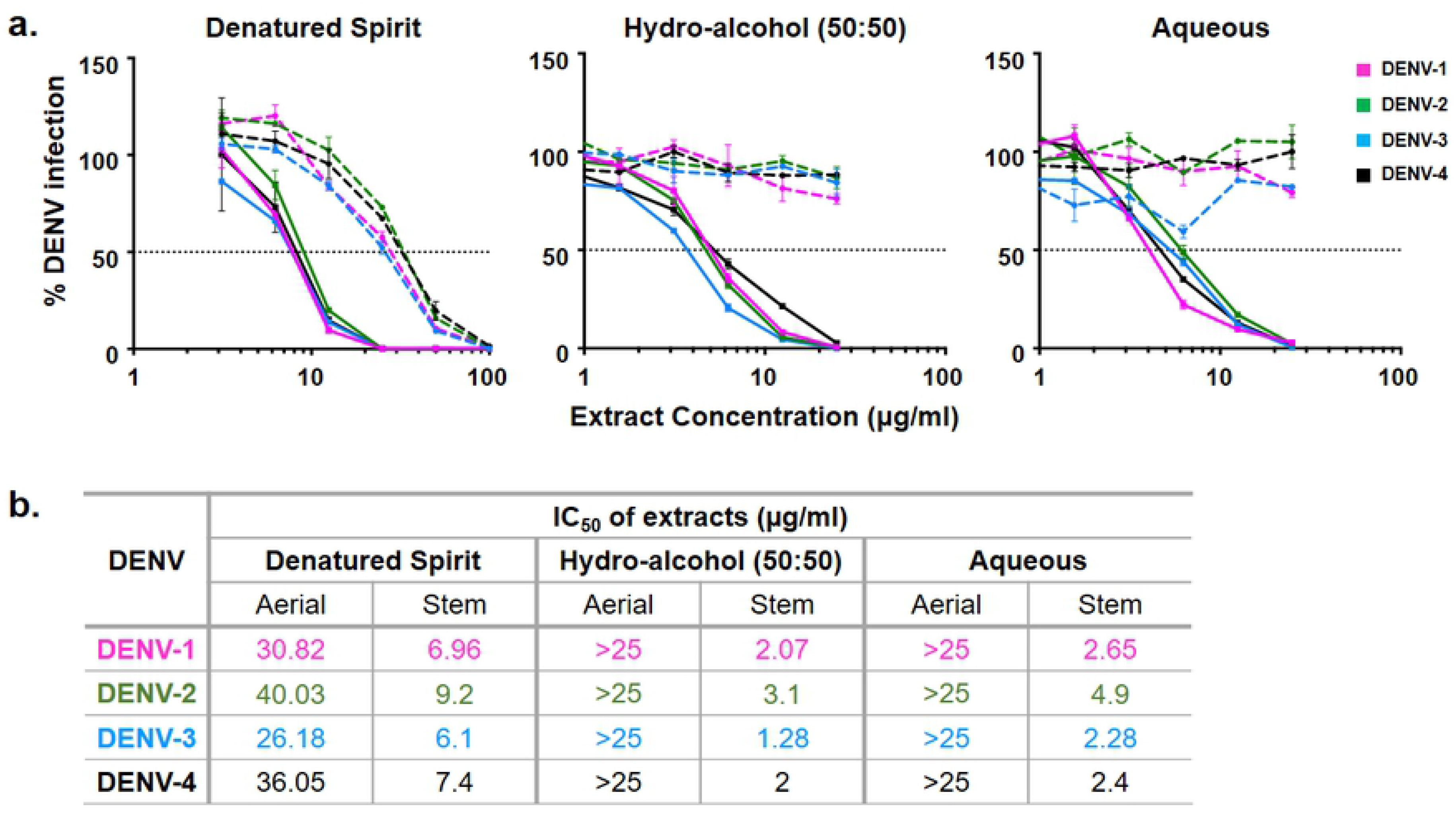
Evaluation of extracts of aerial and stem parts of *C. hirsutus* prepared using various extraction solvents: Extracts of aerial and stem only parts of *C. hirsutus* were prepared in different solvents viz., denatured spirit, hydro-alcohol (50:50) and aqueous. The anti-dengue activity of each of these extracts at various concentrations was evaluated against DENV-1 (magenta curve), DENV-2 (green curve), DENV-3 (blue curve) and DENV-4 (black curve) by flow-cytometry-based virus inhibition assay. (a) The % DENV infection relative to virus control achieved is represented graphically for denatured spirit (left panel), hydro-alcohol, 50:50 (middle panel) and aqueous (right panel) aerial (dashed curves) and stem (solid curves) extracts. (b) The concentration of extract (μg/ml) that resulted in 50% inhibition of viral infection as compared to virus control (represented by horizontal dotted line in panel ‘a’), calculated as IC_50_ using Graphpad Prism, is shown in the table for all the extracts.

### Dose-dependent inhibition of secretion of DENV and its antigen by AQCH

The measurement of anti-DENV activity of AQCH through flow-cytometry-based virus inhibition assay quantitates the cytosolic virus 46 hr post-infection. In order to ascertain the DENV inhibition and its kinetics, we evaluated the impact of AQCH on the secreted virus and its secretory antigen NS1 for up to 6 days post infection (Fig 3).

**Fig 3:**
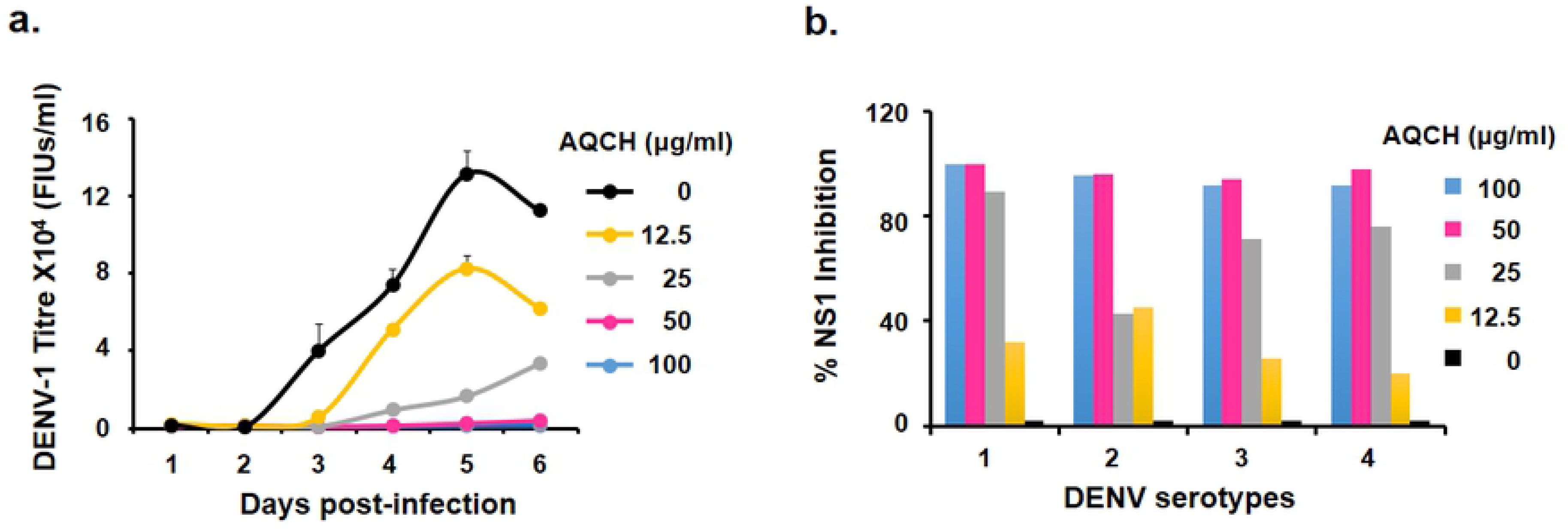
AQCH inhibits secretion of DENV and its antigen, NS1, in a dose-dependent manner: DENV 1-4 infected Vero cells were incubated with 2 fold dilutions of AQCH ranging from 100 to 12.5 μg/ml. Aliquots of the supernatant were collected on each day till day 6 post-infection and analyzed for (a) amount of secreted DENV-1 from days 1-6 through FACS based virus titration assay yielding FIUs/ml, and (b) % inhibition of secretion of viral antigen, NS1, evaluated through commercial ELISA kit on day 6 for all the four DENV serotypes.

Aliquots of culture supernatant from DENV 1-4 infected Vero cells were collected from day 1 to 6 post-infection and were analysed for titration of secreted DENV and NS1 by flow-cytometry based virus inhibition assay and commercial ELISA kit, respectively. It was observed that AQCH at 100 and 50 μg/ml was highly effective in completely inhibiting the secretion of DENV up to 6 days of the experiment and this inhibition was observed to be dose-dependent (Fig 3a; data shown only with DENV-1). Analysis of NS1 levels in the collected supernatants for all the four DENVs on day 6 corroborated this result as 100 and 50 μg/ml of AQCH exhibited 100% inhibition of release of NS1 and this inhibition decreased with the decrease in concentration of AQCH (Fig 3b).

### Chemical fingerprinting of AQCH and identification of chemical markers

Another batch of AQCH (ID: KL/DBE/002/18) was prepared and confirmed for its pan anti-dengue activity by flow-cytometry-based virus inhibition assay (Fig 4a). Additionally, the extent of *in vitro* cytotoxicity caused to Vero cells by AQCH was also evaluated by MTT assay and the CC_50_ was determined to be ~90 μg/ml (Fig 4a). HPLC chromatography was performed on this AQCH batch for its chemical profiling; the chromatogram obtained is shown in Fig 4b. This was followed by isolation of five marker compounds using chromatographic methods, which were characterised using advanced 1D and 2D NMR spectroscopic and mass analysis. Marker compounds were identified to be Sinococuline (1), Magnoflorine (2), 20-Hydroxyecdysone (3), Makisterone-A (4), and Coniferyl alcohol (5) (Fig 4c).

**Fig 4:**
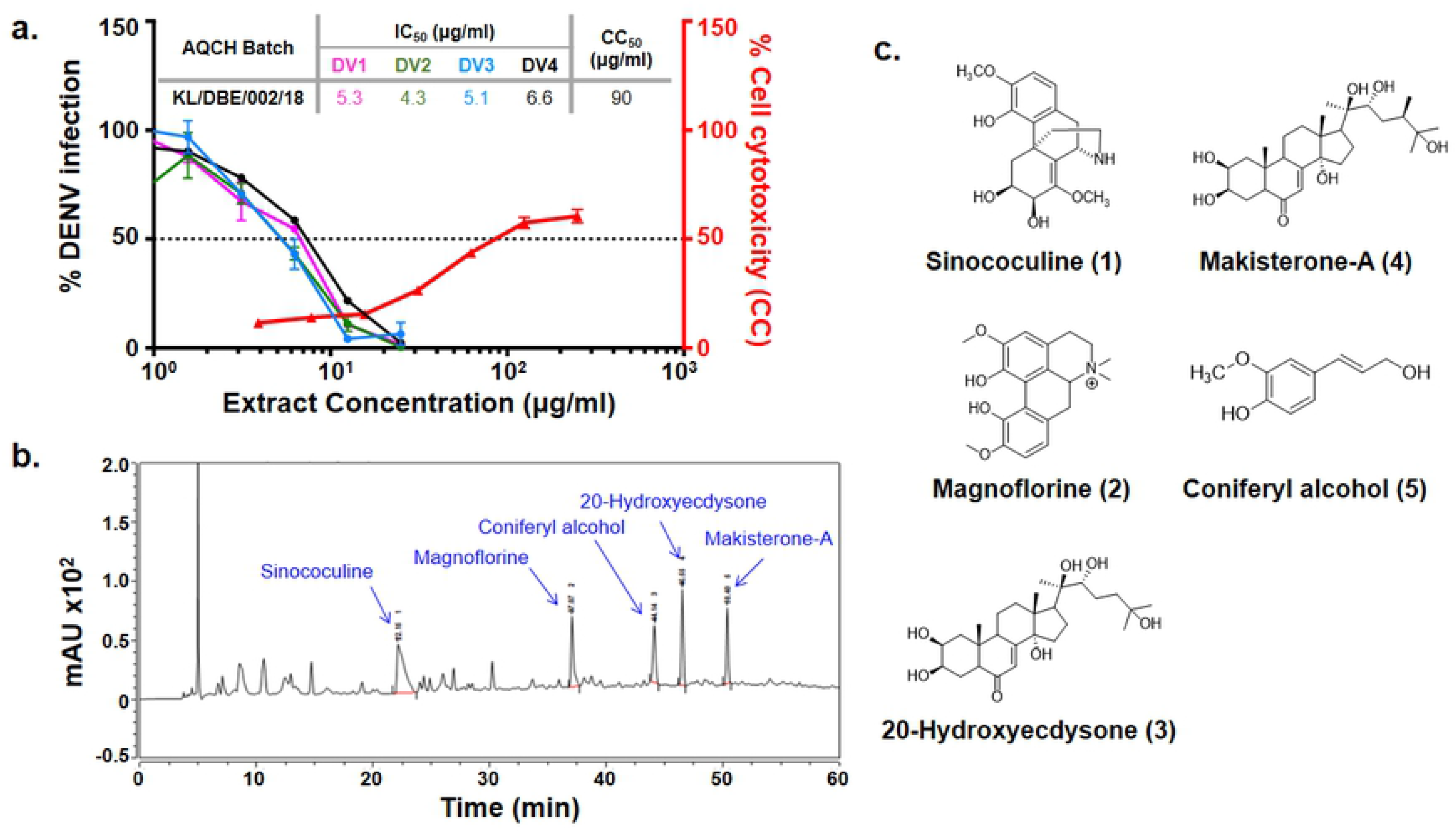
Chemical fingerprinting of AQCH: (a) AQCH batch KL/DBE/002/18 was prepared and its anti-dengue activity against DENV-1 (magenta curve), DENV-2 (green curve), DENV-3 (blue curve) and DENV-4 (black curve) was confirmed by flow-cytometry-based virus inhibition assay as represented by graph of % DENV infection on left y-axis and extract concentration. The extent of cell cytotoxicity caused by AQCH (represented by red curve) was also measured by MTT assay that is reflected on the right y-axis of the graph as % cell cytotoxicity for the given extract concentrations on the x-axis. The CC_50_ and IC_50_ values corresponding to the concentration of AQCH that is toxic for 50% of the cells and at which 50% of DENV infection is inhibited as compared to virus control, respectively has been represented by a dotted horizontal line; a table of IC_50_ and CC_50_ values has been provided as an inset. (b) HPLC chemical fingerprinting profile of AQCH with the peaks corresponding to the five identified marker compounds annotated. (c) Chemical structure of the five marker compounds (1-5).

### Evaluation of robustness and consistency in the preparation of AQCH

Various batches of AQCH were prepared utilising one of the three drying methods-rotary vapour drying, vacuum tray drying and spray drying. Irrespective of the method used for drying, the *in vitro* anti-dengue activity of all the extract batches prepared was comparable (Fig 5a). The HPLC chromatograms of three batches corresponding to the three drying methods were observed to be overlapping (Fig 5b), with high degree of consistency in retention times of the five marker compounds (Fig 5c). This indicates that the AQCH extract preparation method is consistent and robust, and the choice of drying method does not have any implication on its chemical profiling and biological activity. Thus, spray drying was considered as the method of choice, as it resulted in the formation of free-flowing finer extract in a shorter span of time, which is industrially more compatible.

**Fig 5:**
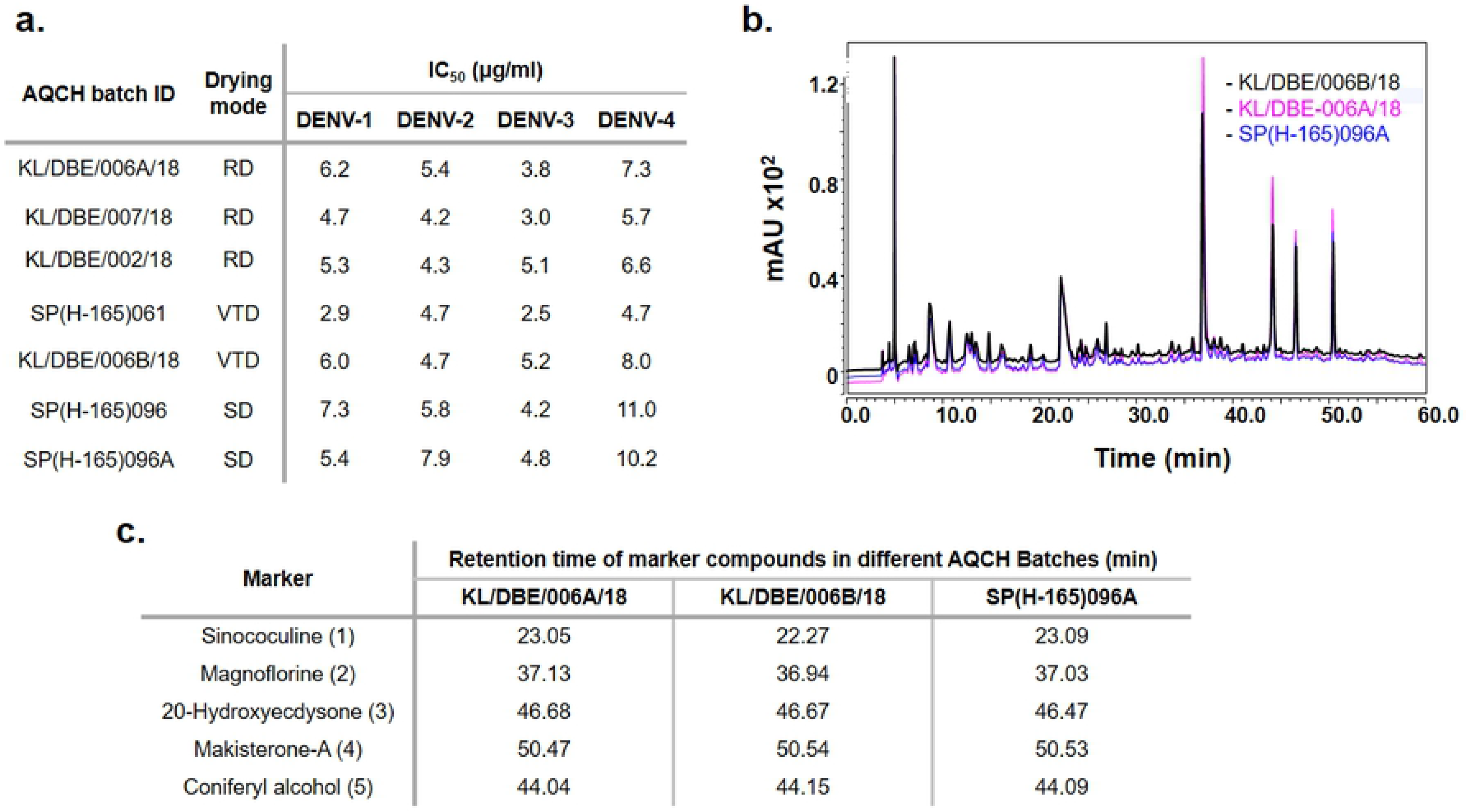
AQCH preparation method is consistent and robust: Various batches of AQCH were prepared and dried through either of the three different methods viz., rotary vapour drying (RD), vacuum tray drying (VTD) or spray drying (SD). The effect of drying method was evaluated through the assessment of (a) anti-dengue activity by flow-cytometry-based virus inhibition assay yielding IC_50_ values (concentration of the extract required to reduce the DENV infection by 50% as compared to virus control), and (b,c) chemical fingerprinting profile; an overlay HPLC chromatograms of the three batches corresponding to the three drying conditions and a table of retention time of five marker compounds are shown in panels ‘b’ and ‘c’, respectively.

### AQCH provides protection against lethal infection of DENV-2 in AG129 mouse model

AG129 are immune-compromised mice deficient in interferon α/β and γ receptor signalling, which allows propagation of mouse adapted DENV-2 S221 strain to result in development of disease. Hence, this mouse model was used to evaluate the efficacy of AQCH *in vivo*. The design of the assay is depicted in Fig 6a, where the AG129 mice were infected through intravenous route with a lethal dose of DENV-2 S221 (1.0 × 10^5^ FIUs). This was followed by oral feeding for 5 days with either 25 mg/kg/dose (Group ‘V + AQCH 25 mg/kg/dose QID’, blue curve) or 8.25 mg/kg/dose (Group ‘V + AQCH 8.25 mg/kg/dose QID’, pink curve) AQCH QID and were monitored for survival, morbidity score and weight change for up to 15 days post-infection. Non-infected and non-AQCH fed (Group ‘Uninfected’, black curve) and non-infected but AQCH fed (Group ‘Only AQCH’, orange curve) AG129 mice groups served as negative controls, while virus infected but not fed with AQCH AG129 mice group (Group ‘V’, grey curve) served as positive control. AG129 mice of Group ‘V’ did not survive beyond six days, and exhibited highest morbidity scores and % body weight change (Fig 6b-d, grey curve). However, infected AG129 mice that were fed with 25 and 8.25 mg/kg/dose QID AQCH were significantly protected (p<0.05), exhibiting 100% and 50% survival, respectively (Fig 6b, blue and pink curves, respectively); their morbidity scores and % body weight change too improved gradually after peaking around day 4-6 (Fig 6c,d). The negative control groups, ‘Uninfected’ and ‘Only AQCH’, did not exhibit any mortality (Fig 6b, black and orange curves, respectively) and there was neither significant morbidity nor reduction in body weight observed (Fig 6c,d; black and orange curves, respectively).

**Fig 6:**
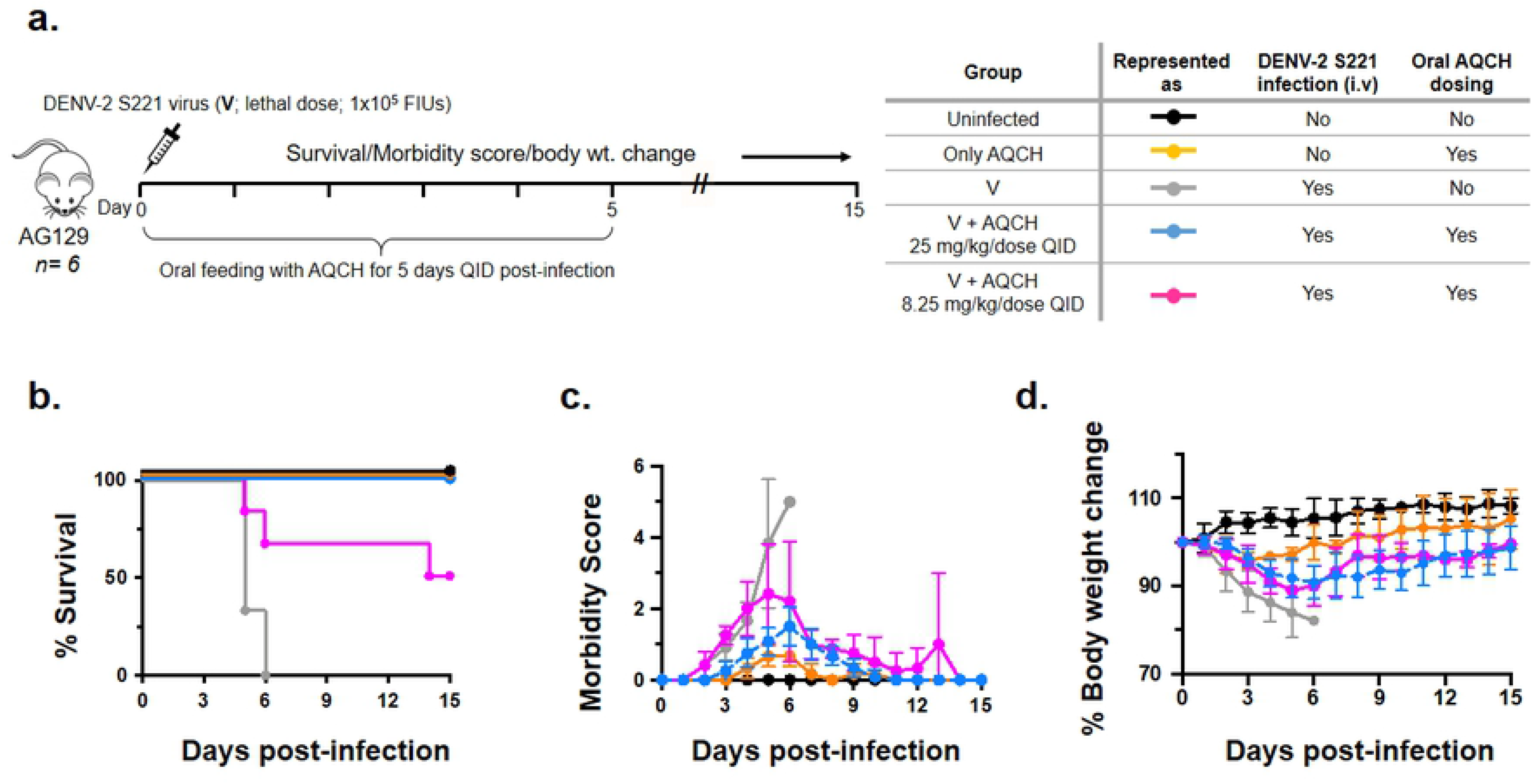
AQCH protects AG129 mice from DENV-2 S221 lethal infection: (a) Schematic representation of the design of experiment using five groups of AG129 mice (n= 6). ‘Uninfected’ group represented by black curve, was neither infected with DENV-2 S221 nor dosed with AQCH. ‘Only AQCH’ group, represented by orange curve, was not infected with DENV-2 S221 but received AQCH dose (25 mg/kg/dose QID). ‘V’ group, represented by grey curve, was infected with DENV-2 S221 but was not dosed with AQCH. Mice in the remaining two groups were infected with DENV-2 S221 and were dosed either with 25 mg/kg/dose, QID (blue curve) or 8.25 mg/kg/dose, QID (pink curve). DENV-2 S221 infection was given i.v. at a lethal dose of 1.0 × 10^5^ FIUs, while AQCH was dosed orally post-infection. All the groups were monitored for (b) survival, (c) morbidity score, and (d) body weight change over the next 15 days post-infection. Survival data (panel ‘b’) were analysed by Log-Rank (Mantel-Cox) test for statistical evaluation of level of significance in difference in survival rates. Survival of mice in ‘V+AQCH 25 mg/kg/dose QID’ and ‘V+AQCH 8.25 mg/kg/dose QID’ groups was not significantly different from each other (p= 0.14), but differed significantly from Group ‘V’ survival (p= 0.006 and p= 0.016, respectively). The p value <0.05 was considered significant. The Morbidity score in panel ‘c’ was based on 5 point system: 0.5, mild ruffled fur; 1.0, ruffled fur; 1.5, compromised eyes; 2, compromised eyes with hunched back; 2.5, loose stools; 3.0, limited movement; 3.5, no movement/hind leg paralysis; 4.0, euthanized if cumulative score was 5. Body weight in panel ‘d’ was monitored twice a day in the morning and evening, and the mean taken for plotting the graph.

### Feasibility of clinical evaluation of AQCH

With the exhibition of *in vitro* and *in vivo* anti-dengue potency by AQCH, it became evident that it has a strong potential to be developed as a drug, however, its clinical suitability was yet to be evaluated. Our first study on that front was to evaluate its interaction with paracetamol which is a standard-of-care drug for treating dengue fever. In this study, a range of concentration of AQCH was separately evaluated against DENV-1 through flow-cytometry-based virus inhibition assay in absence (0 μg/ml) and presence of 1, 10 and 100 μg/ml paracetamol. Importantly, the anti-dengue activity of AQCH was found to be unaffected by paracetamol in this experiment (Fig 7).

**Fig 7:**
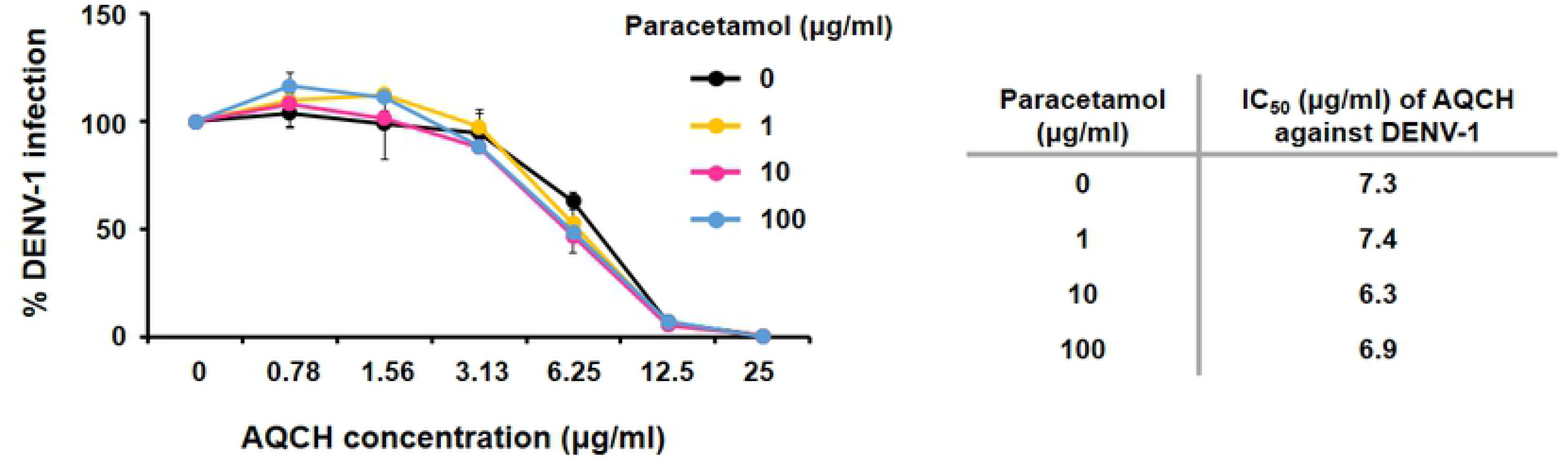
Paracetamol does not inhibit the anti-dengue activity of AQCH. DENV-1 infected Vero cells were treated with various concentrations of AQCH (0-25 μg/ml) in absence (black curve) and presence of 1 (orange curve), 10 (magenta curve) and 100 (blue curve) μg/ml of paracetamol separately. The % DENV-1 infection achieved under these conditions was evaluated in a flow-cytometry-based virus inhibition assay, which is depicted in the graph on the left panel. Concentration of AQCH that led to 50% reduction in DENV-1 infection as compared to virus control was calculated separately for each condition as its corresponding IC_50_ and is depicted in the table on the right panel.

The next aspect of AQCH evaluation was its tablet formulation for clinical utility. Thus, 100, 300 and 500 mg strengths of AQCH tablets were formulated and subjected to accelerated and long term stability studies along with the AQCH extract batch from which the tablets were formulated. Samples from stability study were analysed for *in vitro* anti-DENV-2 activity by flow-cytometry-based virus inhibition assay to evaluate any deterioration or loss in bioactivity upon storage under the conditions tested. It was found that there was no significant change in the anti-DENV-2 activity (Table 1) of AQCH and AQCH tablets of all the three strengths (100, 300 and 500 mg) under the conditions tested up to 6 months. The long-term stability study is on-going and samples will be evaluated up to 3 years of storage. This data was encouraging as it ensured the feasibility of formulating AQCH into a stable tablet dosage form, which is advantageous for its clinical evaluation.

**Table 1:** Anti-DENV-2 activity of AQCH and AQCH tablets under the stated conditions of storage.

| AQCH Extract/<br>Tablet (Batch ID) | Storage condition | DENV-2 IC<sub>50</sub> (µg/ml)<sup>a</sup> |  |  |  |
| --- | --- | --- | --- | --- | --- |
|  |  | 1 Month | 2 Month | 3 Month | 6 Month |
| **Extract**<br>**(FCH1901002)** | 40±2°C, 75±5% RH | 4.2 | 4.0 | 5.4 | 7.2 |
|  | 30±2°C, 65±5% RH | nd* | nd* | 6.1 | 5.7 |
| **Tablet 100 mg**<br>**{RYP(6665)079A}** | 40±2°C, 75±5% RH | 4.7 | 4.6 | 7.3 | 5.4 |
|  | 30±2°C, 65±5% RH | nd* | nd* | 7.8 | 4.7 |
| **Tablet 300 mg**<br>**{RYP(6665)079B}** | 40±2°C, 75±5% RH | 5.2 | 6.2 | 6.2 | 5.3 |
|  | 30±2°C, 65±5% RH | nd* | nd* | 6.1 | 4.0 |
| **Tablet 500 mg**<br>**{RYP(6665)079C}** | 40±2°C, 75±5% RH | 4.6 | 2.7 | 5.1 | 5.1 |
|  | 30±2°C, 65±5% RH | nd* | nd* | 6.8 | 4.7 |

<sup>a</sup> Anti-DENV-2 activity is measured as IC<sub>50</sub>, which corresponds to the concentration at which 50% of the virus is inhibited with respect to virus control
\*nd: not determined; these specific storage conditions were for long-term stability studies and were therefore not sampled on 1<sup>st</sup> and 2<sup>nd</sup> month of storage

## Discussion

Plants have been traditionally and historically used worldwide for their therapeutic potential since time immemorial. Their therapeutic usefulness has been documented all around the globe in various traditional literatures. Hence, these classical troves which detail medicinal utility of plants are an attractive repertoire of knowledge that could be explored through contemporary methods for the development of safe and effective therapies for various maladies. This has ushered the development of many plant-derived molecules which today are in clinical use globally, morphine being the first FDA-approved plant derived molecule [35]. We referred to one of the world’s oldest holistic healing system, the Indian traditional medicine of Ayurveda, in our quest to develop an effective therapy against dengue.

Dengue is one of the world’s rapidly spreading arboviral diseases with the incidence of symptomatic dengue doubling every decade [36]. The highest burden of this disease lies in Southeast Asia with India being one of the epicentres [7, 36]. According to a study, actual dengue cases in India are ~282 times higher than that reported annually, having an economic impact of USD ~1.11 billion [37]. Thus, there is a dire need of an effective dengue vaccine and/or drug to fight against dengue. Dengvaxia is the world’s first approved dengue vaccine, however, its utility is limited to only seropositive adults in dengue-endemic nations due to concerns of vaccine-induced enhancement of virus infection [38]. In parallel, rigorous efforts are also being made towards dengue therapeutics with the evaluation of novel and repurposed drugs against DENV [39,40]. However, none of the drugs have yet succeeded in proof-of-concept trials and thus, a dengue antiviral still remains an unmet need.

Guided by the Indian Ayurveda literature, our group had earlier evaluated 19 medicinal plants for their anti-dengue activity that led to the identification of *C. pareira* of Menispermaceae family as the most potent plant [28]. Our continued exploration of more plants belonging to Menispermaceae family through scientific and Ayurvedic literature [32–34] led us to select *C. hirsutus* for the evaluation of its anti-dengue activity. Thus, methanolic extracts of aerial parts of *C. hirsutus* and *C. pareira* were compared head-on in an *in vitro* flow-cytometry-based virus inhibition assay. It was observed that *C. hirsutus* is a significantly more potent anti-dengue herb than *C. pareira* (Fig 1).

For greater regulatory acceptance, we wanted to evaluate if methanol could be replaced with milder solvents like denatured spirit, hydro-alcohol (50:50) or water for extract preparation. Thus, each of these solvents were explored for separate extract preparations of stem and aerial parts of *C. hirsutus*, and evaluated for their anti-dengue activity (Fig 2). This experiment yielded two important outcomes. First, irrespective of the solvent used, stem part of *C. hirsutus* was significantly more potent in its anti-dengue activity than the aerial part. Second, stem extract prepared in water was comparable in its anti-dengue activity to other tested solvents. Thus, aqueous extract of stem of *C. hirsutus* (AQCH) was selected as the extract of choice for further evaluations owing to greater regulatory acceptance of water as a solvent.

The effect of AQCH on the secretion of DENV and its secretory antigen NS1 was monitored over a period of 6 days through *in vitro* evaluations (Fig 3). AQCH was found to inhibit the secretion of both DENV and NS1 in a dose-dependent manner, with complete inhibition being observed at 100 and 50 μg/ml extract. This is relevant because DENV load and NS1 have been implicated in dengue disease pathogenesis in humans [41,42].

The cytotoxicity of AQCH was determined *in vitro* on Vero cells through MTT assay and the CC_50_ was observed to be more than 10-fold higher as compared to its IC_50_ (Fig 4a), indicating a good therapeutic window for AQCH. We further evaluated the industrial viability of AQCH production. For this an AQCH batch was profiled through HPLC chromatography and five marker compounds-Sinococuline, Magnoflorine, 20-Hydroxyecdysone, Makisterone-A and Coniferyl alcohol were identified (Fig 4b,c). The chemical profiling data of a bioactive batches of AQCH were used to monitor the quality of extract prepared during optimization of extraction method. Thus, various batches of AQCH were prepared using three different drying methods viz. rotary vapour, vacuum tray or spray drying. All the AQCH batches were found to exhibit similar anti-dengue activity (Fig 5a) irrespective of the drying process, indicating the robustness and consistency of the method of extraction. This was corroborated by their HPLC chemical fingerprinting that yielded similar chromatograms (Fig 5b,c). Spray-dried AQCH extracts were utilized for further evaluations due to greater industrial compatibility.

AQCH was evaluated for its protective efficacy *in vivo* in the AG129 mouse model (Fig 6), which is an established model for the evaluation of antivirals [43,44]. AG129 mice being deficient in IFN α/β and γ receptors allow propagation of DENV and development of dengue disease-like symptoms [45]. AQCH, when fed at 25 mg/kg/dose QID, was found to provide 100% protection to AG129 mice that were lethally infected with DENV-2; 8.25 mg/kg/dose QID AQCH resulted in 50% protection (Fig 6). Demonstration of potent anti-dengue activity by AQCH in *in vitro* and *in vivo* analyses lays the ground for its clinical development. Paracetamol, a standard care drug in treating dengue, was found to have no effect on the anti-dengue activity of AQCH (Fig 7). Also, spray-dried AQCH was formulated into tablets of various strengths, which were found to be stable upon storage (Table 1). This supports the case for the clinical use of AQCH tablets along with paracetamol in treating dengue.

In conclusion, this is the first study reporting an aqueous extract of the stem of *C. hirsutus* to possess significant pan anti-dengue activity; the extraction process is robust and consistent, making this plant industrially viable for further clinical development. At this time when there is no approved anti-dengue drug available, this phytopharmaceutical formulation can be a breakthrough in providing a safe and effective drug against dengue, which is urgently needed globally.

## Acknowledgments

Authors are thankful to Dr. Mohan Prasad, Dr. Azadar Khan, Mr. Narendra Lakkad, Dr. Romi Singh, Dr. Atul Raut, Mr. Gaurav Sahal, Mr. Rakesh Sinha, Mr. Kohinoor Das from Sun Pharmaceutical Industries Ltd., Dr. Dinakar Salunke from International Centre for Genetic Engineering and Biotechnology, New Delhi, and Dr. Mohammad Aslam from Department of Biotechnology, Government of India for their support during the entire study period.

## Author Contributions

**Conceptualisation** NK and AAL

**Project Administration:** RK

**Data Generation: BRM collection-** SP, KN, SG; **Extract preparation-** SP, KN; **Virus inhibition assays-** AP, HB, RKS; **NS1 assay-** AP; **MTT Assay-** RKR; **Animal experiments-** RS; **Tablet formulation and stability studies-** TJ, BV, RP, HM, SM; **Chemical fingerprinting and marker compound isolation-** DA, VS, PG, APG, DS, YSB, RV

**Data Analysis:** AP, RS, HB, UA, RKR, NK, RSo, AAL, DA, VS, PG, APG, DS, YSB, RV, SP, KN, RP, HM, SM, TJ, BV, RK

**Data Curation:** UA

**Manuscript Writing:** RKR, UA

**Manuscript Review:** NK, AAL, UA, RKR

## References

1) WHO weblink on dengue control. What is Dengue? Available at https://www.who.int/denguecontrol/disease/en/. Accessed on 13th June 2020.

2) Gubler DJ. Dengue, Urbanization and Globalization: The Unholy Trinity of the 21(st) Century. Trop Med Health. 2011;39(4 Suppl):3–11.

3) Struchiner CJ, Rocklov J, Wilder-Smith A, Massad E. Increasing Dengue Incidence in Singapore over the Past 40 Years: Population Growth, Climate and Mobility. PLoS One. 2015;10(8):e0136286.

4) Wilder-Smith A, Ooi EE, Horstick O, Wills B. Dengue. Lancet. 2019;393(10169):350–63.

5) Andrade EH, Figueiredo LB, Vilela AP, Rosa JC, Oliveira JG, Zibaoui HM, et al. Spatial-Temporal Co-Circulation of Dengue Virus 1, 2, 3, and 4 Associated with Coinfection Cases in a Hyperendemic Area of Brazil: A 4-Week Survey. Am J Trop Med Hyg. 2016;94(5):1080–4.

6) Shrivastava S, Tiraki D, Diwan A, Lalwani SK, Modak M, Mishra AC, et al. Co-circulation of all the four dengue virus serotypes and detection of a novel clade of DENV-4 (genotype I) virus in Pune, India during 2016 season. PLoS One. 2018;13(2):e0192672.

7) Bhatt S, Gething PW, Brady OJ, Messina JP, Farlow AW, Moyes CL, et al. The global distribution and burden of dengue. Nature. 2013;496(7446):504–7.

8) Brady OJ, Gething PW, Bhatt S, Messina JP, Brownstein JS, Hoen AG, et al. Refining the global spatial limits of dengue virus transmission by evidence-based consensus. PLoS Negl Trop Dis. 2012;6(8):e1760.

9) Murhekar MV, Kamaraj P, Kumar MS, Khan SA, Allam RR, Barde P, et al. Burden of dengue infection in India, 2017: a cross-sectional population based serosurvey. Lancet Glob Health. 2019;7(8):e1065–e73.

10) Shepard DS, Undurraga EA, Halasa YA, Stanaway JD. The global economic burden of dengue: a systematic analysis. Lancet Infect Dis. 2016;16(8):935–41.

11) Dengue: Guidelines for Diagnosis, Treatment, Prevention and Control: New Edition. WHO Guidelines Approved by the Guidelines Review Committee. Geneva2009.

12) Muller DA, Depelsenaire AC, Young PR. Clinical and Laboratory Diagnosis of Dengue Virus Infection. J Infect Dis. 2017;215(suppl_2):S89–S95.

13) Aguiar M, Stollenwerk N, Halstead SB. The risks behind Dengvaxia recommendation. Lancet Infect Dis. 2016;16(8):882–3.

14) World Health O. Dengue vaccine: WHO position paper, July 2016 - recommendations. Vaccine. 2017;35(9):1200–1.

15) Thomas SJ, Yoon IK. A review of Dengvaxia(R): development to deployment. Hum Vaccin Immunother. 2019;15(10):2295–314.

16) Beesetti H, Khanna N, Swaminathan S. Investigational drugs in early development for treating dengue infection. Expert Opin Investig Drugs. 2016;25(9):1059–69.

17) Lim SP. Dengue drug discovery: Progress, challenges and outlook. Antiviral Res. 2019;163:156–78.

18) Dighe SN, Ekwudu O, Dua K, Chellappan DK, Katavic PL, Collet TA. Recent update on anti-dengue drug discovery. Eur J Med Chem. 2019;176:431–55.

19) Tricou V, Minh NN, Van TP, Lee SJ, Farrar J, Wills B, et al. A randomized controlled trial of chloroquine for the treatment of dengue in Vietnamese adults. PLoS Negl Trop Dis. 2010;4(8):e785.

20) Low JG, Sung C, Wijaya L, Wei Y, Rathore APS, Watanabe S, et al. Efficacy and safety of celgosivir in patients with dengue fever (CELADEN): a phase 1b, randomised, double-blind, placebo-controlled, proof-of-concept trial. Lancet Infect Dis. 2014;14(8):706–15.

21) Whitehorn J, Nguyen CVV, Khanh LP, Kien DTH, Quyen NTH, Tran NTT, et al. Lovastatin for the Treatment of Adult Patients With Dengue: A Randomized, Double-Blind, Placebo-Controlled Trial. Clin Infect Dis. 2016;62(4):468–76.

22) Nguyen NM, Tran CN, Phung LK, Duong KT, Huynh Hle A, Farrar J, et al. A randomized, double-blind placebo controlled trial of balapiravir, a polymerase inhibitor, in adult dengue patients. J Infect Dis. 2013;207(9):1442–50.

23) Tam DT, Ngoc TV, Tien NT, Kieu NT, Thuy TT, Thanh LT, et al. Effects of short-course oral corticosteroid therapy in early dengue infection in Vietnamese patients: a randomized, placebo-controlled trial. Clin Infect Dis. 2012;55(9):1216–24.

24) Singh PK, Rawat P. Evolving herbal formulations in management of dengue fever. J Ayurveda Integr Med. 2017;8(3):207–10.

25) Parida MM, Upadhyay C, Pandya G, Jana AM. Inhibitory potential of neem (Azadirachta indica Juss) leaves on dengue virus type-2 replication. J Ethnopharmacol. 2002;79(2):273–8.

26) Jain M, Ganju L, Katiyal A, Padwad Y, Mishra KP, Chanda S, et al. Effect of Hippophae rhamnoides leaf extract against Dengue virus infection in human blood-derived macrophages. Phytomedicine. 2008;15(10):793–9.

27) Kasture PN, Nagabhushan KH, Kumar A. A Multi-centric, Double-blind, Placebo-controlled, Randomized, Prospective Study to Evaluate the Efficacy and Safety of Carica papaya Leaf Extract, as Empirical Therapy for Thrombocytopenia associated with Dengue Fever. J Assoc Physicians India. 2016;64(6):15–20.

28) Sood R, Raut R, Tyagi P, Pareek PK, Barman TK, Singhal S, et al. Cissampelos pareira Linn: Natural Source of Potent Antiviral Activity against All Four Dengue Virus Serotypes. PLoS Negl Trop Dis. 2015;9(12):e0004255.

29) Marya BH, Bothara SB. Ethnopharmacological properties of Cocculus hirsutus (L.) Diels-A review. International Journal of Pharmaceutical Sciences Review and Research. Volume 7, Issue 1, March – April 2011; Article-022

30) Ganapaty S, Dash GK, Subburaju T, Suresh P. Diuretic, laxative and toxicity studies of Cocculus hirsutus aerial parts. Fitoterapia. 2002;73(1):28–31.

31) Lambeth CR, White LJ, Johnston RE, de Silva AM. Flow cytometry-based assay for titrating dengue virus. J Clin Microbiol. 2005;43(7):3267–72.

32) Barbosa-Filho JM, Da-Cunha EVL, Gray AI. Alkaloids of the Menispermaceae. In: The Alkaloids: Chemistry and Biology. Academic Press; 2000; 54: 1–190.

33) Hishiki T, Kato F, Tajima S, Toume K, Umezaki M, Takasaki T, Miura T. Hirsutine, an indole alkaloid of Uncaria rhynchophylla, inhibits late step in dengue virus life cycle. Front Microbiol. 2017: 8:1674.doi: 10.3389/fmicb.2017.01674

34) Rasheed T, Khan MNI, Zhadi SSA. Hirsutine: A new alkaloid from Cocculus hirsutus. J Nat Prod. 1991; 54(2): 582–584.

35) Li FS, Weng JK. Demystifying traditional herbal medicine with modern approach. Nat Plants. 2017;3:17109.

36) Stanaway JD, Shepard DS, Undurraga EA, Halasa YA, Coffeng LE, Brady OJ, et al. The global burden of dengue: an analysis from the Global Burden of Disease Study 2013. Lancet Infect Dis. 2016;16(6):712–23.

37) Shepard DS, Halasa YA, Tyagi BK, Adhish SV, Nandan D, Karthiga KS, et al. Economic and disease burden of dengue illness in India. Am J Trop Med Hyg. 2014;91(6):1235–42.

38) Halstead SB. Dengvaxia sensitizes seronegatives to vaccine enhanced disease regardless of age. Vaccine. 2017;35(47):6355–8.

39) Botta L, Rivara M, Zuliani V, Radi M. Drug repurposing approaches to fight Dengue virus infection and related diseases. Front Biosci (Landmark Ed). 2018;23:997–1019.

40) Low JG, Ooi EE, Vasudevan SG. Current Status of Dengue Therapeutics Research and Development. J Infect Dis. 2017;215(suppl_2):S96–S102.

41) Chen HR, Lai YC, Yeh TM. Dengue virus non-structural protein 1: a pathogenic factor, therapeutic target, and vaccine candidate. J Biomed Sci. 2018;25(1):58.

42) Tricou V, Minh NN, Farrar J, Tran HT, Simmons CP. Kinetics of viremia and NS1 antigenemia are shaped by immune status and virus serotype in adults with dengue. PLoS Negl Trop Dis. 2011;5(9):e1309.

43) Chan KW, Watanabe S, Kavishna R, Alonso S, Vasudevan SG. Animal models for studying dengue pathogenesis and therapy. Antiviral Res. 2015;123:5–14.

44) Watanabe S, Vasudevan SG. Evaluation of dengue antiviral candidates in vivo in mouse model. Methods Mol Biol. 2014;1138:391–400.

45) Julander JG, Perry ST, Shresta S. Important advances in the field of anti-dengue virus research. Antivir Chem Chemother. 2011;21(3):105–16.

